# Maternal vgll4a promotes blastoderm cohesion enabling yap1-mediated mechano-transduction during zebrafish epiboly

**DOI:** 10.1101/2020.12.01.407478

**Authors:** Carlos Camacho-Macorra, Noemí Tabanera, Paola Bovolenta, Marcos J Cardozo

## Abstract

Cellular cohesion provides tissue tension, which is then sensed by the cytoskeleton and decoded by the activity of mechano-transducers, such as the transcriptional cofactor Yap1, thereby enabling morphogenetic responses in multi-cellular organisms. How cell cohesion is regulated is nevertheless unclear. Here we show that, zebrafish epiboly progression, a prototypic morphogenetic event that depends on Yap activity, requires the maternal contribution of the proposed yap1 competitor vgll4a. In embryos lacking maternal/zygotic *vgll4a* (*MZvgll4a*), spreading epithelial cells are ruffled, blastopore closure is delayed and the expression of the yap1-mediator *arhgap18* is decreased, impairing the actomyosin ring at the syncytial layer. Furthermore, rather than competing with Yap1, vgll4a coordinate the levels of the E-Cadherin/β-catenin adhesion complex components at the blastomere plasma membrane and hence their actin cortex distribution. Taking these results together, we propose that maternal vgll4a may act at epiboly initiation to coordinate blastomere adhesion/cohesion, which is a fundamental piece of the self-sustained bio-mechanical regulatory loop underlying morphogenetic rearrangements during gastrulation.

## Introduction

The three-dimensional organization of multicellular organisms requires morphogenetic tissue rearrangements, which are particularly evident during embryonic development or in pathological conditions, such as wound healing or cancer. These rearrangements are largely coordinated by the interplay among cell-cell interaction, mechanical forces, and signaling pathways, derived, in part, from the surrounding environment. The adaptive morphogenetic outcome critically depends on how the cells integrate and interpret this signal complexity and transmit the message to their nuclei, eliciting a consequent transcriptional response (Wozniak & Chen, 2009).

In a number of organisms, including the teleost zebrafish, epiboly is the first morphogenetic rearrangement of gastrulating embryos (Bruce & Heisenberg, 2020). These cellular rearrangements are relatively simple and, thus, have been often used to understand basic principles of morphogenesis (Bruce, 2016). Indeed, in zebrafish the blastoderm, positioned at the animal pole, is formed by a single layer of loosely packed epithelial cells, known as deep cell layer (DEL), which is covered by the so called epithelial enveloping layer (EVL). The blastoderm interfaces with an underlying epithelium, the yolk syncytial layer (YSL), itself a derivative of the blastoderm marginal cells. During epiboly, the blastoderm thins and moves toward the vegetal pole together with the external YSL (E-YSL). These epithelial spreading and thinning are driven, in part, by a circumferential actomyosin network formed by the E-YSL at its border with the EVL. The contraction of this actomyosin ring pulls the EVL to surround the yolk, further dragging the DEL toward the vegetal pole (Bruce, 2016). This process culminates at the end of gastrulation when the blastoderm seals around the yolk in a process known as blastopore closure (Bruce, 2016).

Different studies have shown that several morphogenetic events, including teleost epiboly, involve the function of the transcriptional co-activator Yes1 Associated Transcriptional Regulator (Yap1) and of the highly related TAZ. Yap1 was first described as a downstream nuclear effector of the Hippo signaling pathway, which is involved in the control of cell proliferation (Piccolo *et al*, 2014). However, there is evidence that Yap1 can operate, perhaps independently of Hippo signaling, as a mechano-transducer that links mechanical forces to cellular response (Totaro *et al*, 2018). Cells constantly probe the forces generated in their extracellular environment through plasma membrane adhesive proteins and internal tension adjustments executed by modifications of their actomyosin network. Upon increasing intracellular tension, Yap1, which is normally retained in the cytoplasm, translocates to the nucleus, where it binds to TEA domain (TEAD)-containing transcription factors (TEAD1–TEAD4). The derived complex activates the transcription of target genes, which include F-acting regulators and cell adhesion components, thereby promoting a positive feedback loop that enables cells to counterbalance the external mechanical forces to which they are exposed (Panciera *et al*, 2017).

In this context, recent studies have taken advantage of a *yap1* medaka (*Oryzias latipes*) mutant called *hirame (hir*), to show that *yap1* is required for epiboly progression by regulating the expression of *arhgap18*, a gene encoding a Rho GTPase activating protein involved in the contraction of the actomyosin ring at the E-YSL/EVL interface (Porazinski *et al*, 2015). What are the factors that modulate yap1 function during epiboly remained however without an answer. Here, we have asked if vestigial-like protein 4 (Vgll4) could be one of such factors.

Vgll4 is a transcriptional cofactor that can bind directly to TEAD proteins, via its tandem Tondu domain (TDU), thereby competing for the Yap-TEAD interaction and thus mechanistically acting as an inhibitor of Yap activity (Jiao *et al*, 2014; Koontz *et al*, 2013; Zhang *et al*, 2014). In *Xenopus* and zebrafish the transcripts of the different *vgll4* paralogs show a dynamic developmental expression pattern in different tissues (Barrionuevo *et al*, 2014; Faucheux *et al*, 2010; Xue *et al*, 2018) and in zebrafish the mRNA of two (*vgll4a; vgll4b*) of the three paralogs (*vgll4a, vgll4b* and *vgll4l*) have been detected at the one cell stage (Xue *et al*, 2018), suggesting a possible early function. Although there are additional and species-specific phenotypes that affects later developing organs (Fillatre *et al*, 2019; Wang *et al*, 2020; Xue *et al*, 2019), loss of function studies in mice and analysis of genetic variants in the teleost *Scatophagus argus* suggest that vgll4 participate in the regulation of body size (Yu *et al*, 2019; Feng *et al*, 2019; Yang *et al*, 2020; Suo *et al*, 2020), a trait that is often altered as the consequence of an abnormal gastrulation.

By generating mutants for the three *vgll4* paralogs in zebrafish, we demonstrate that instead of competing with yap1, as possibly expected, maternal but not zygotic vgll4a is required to sustain yap1 signaling, promoting blastomere cohesion and actomyosin contraction driving epiboly progression and normal zebrafish development.

## Results

### *vgll4a* function is required for proper epiboly progression

To determine whether *vgll4* genes contribute to zebrafish epiboly, we generated zebrafish lines in which *vgll4a, vgll4b* and *vgll4l* genes were mutated using CRISPR-Cas9 technology (Fig. S1a). Founders carrying frame-shift mutations for either one of the three *vgll4* paralogs were selected in the F1 generation in order to obtain loss of function lines for each one of the existing Vgll4 proteins (Fig. S1b). Homozygous mutants for *vgll4a, vgll4b* and *vgll4l* genes were obtained in F3. Their offspring was viable with no evident developmental phenotypes (Fig. S1c and data not shown). Similar results were obtained when adult *vgll4a*/*vgll4b* double mutants were analyzed (Fig. S1c).

*vgll4a* and *vgll4b* but not *vgll4l* transcripts are maternally expressed (Xue *et al*, 2018). We thus searched a possible phenotype in the F4 generation, in which *vgll4a* or *vgll4b* parental contribution will be no longer active. By intercrossing adults F3 *vgll4a*, *vgll4b*, *vgll4l* and *vgll4a*/*vgll4b* double mutant, we generated F4 embryos lacking both maternal and zygotic contribution of *vgll4a* (*MZvgll4a*), *vgll4b* (*MZvgll4b*), *vgll4l (MZvgll4l*) and *vgll4a;vgll4b (MZvgll4a;MZvgll4b*). The fertilization rate of the obtained heterozygous *Mvgll4a* and *Mvgll4b* embryos was significantly lower than that of wild type (wt), heterozygous *vgll4a* or heterozygous *vgll4b* embryos (Fig. S2 a-c). In contrast, the fertilization rate of *Mvgll4l* heterozygous was similar to that of wt and heterozygous *vgll4l* embryos (Fig. S2d). This observation suggested that *vgll4a* and *vgll4b* maternal contribution could be important for oogenesis.

Besides the observed decrease in fertilization rate, viable *MZvgll4a* embryos appeared to progress through epiboly at a slower pace, with a delayed blastopore closure than wt embryos (Fig. 1a). To determine if the additional deficiency in maternal *vgll4b* contribution could enhance the phenotype of *MZvgll4a* embryos, we analyzed *MZvgll4a;MZvgll4b* double mutants, which however were no different from *MZvgll4a* embryos (Fig. 1a). Zebrafish mutants with slow epibolic movements present a decreased body length at larva stages (Sun *et al*, 2017). Consistent with this observation, the body length of *MZvgll4a, MZvgll4b* and *MZvgll4a;MZvgll4b* larvae at 3dpf was shorter than those of wt, whereas that of *MZvgll4l* larvae was very comparable (Fig. 1b).

**Figure 1.**
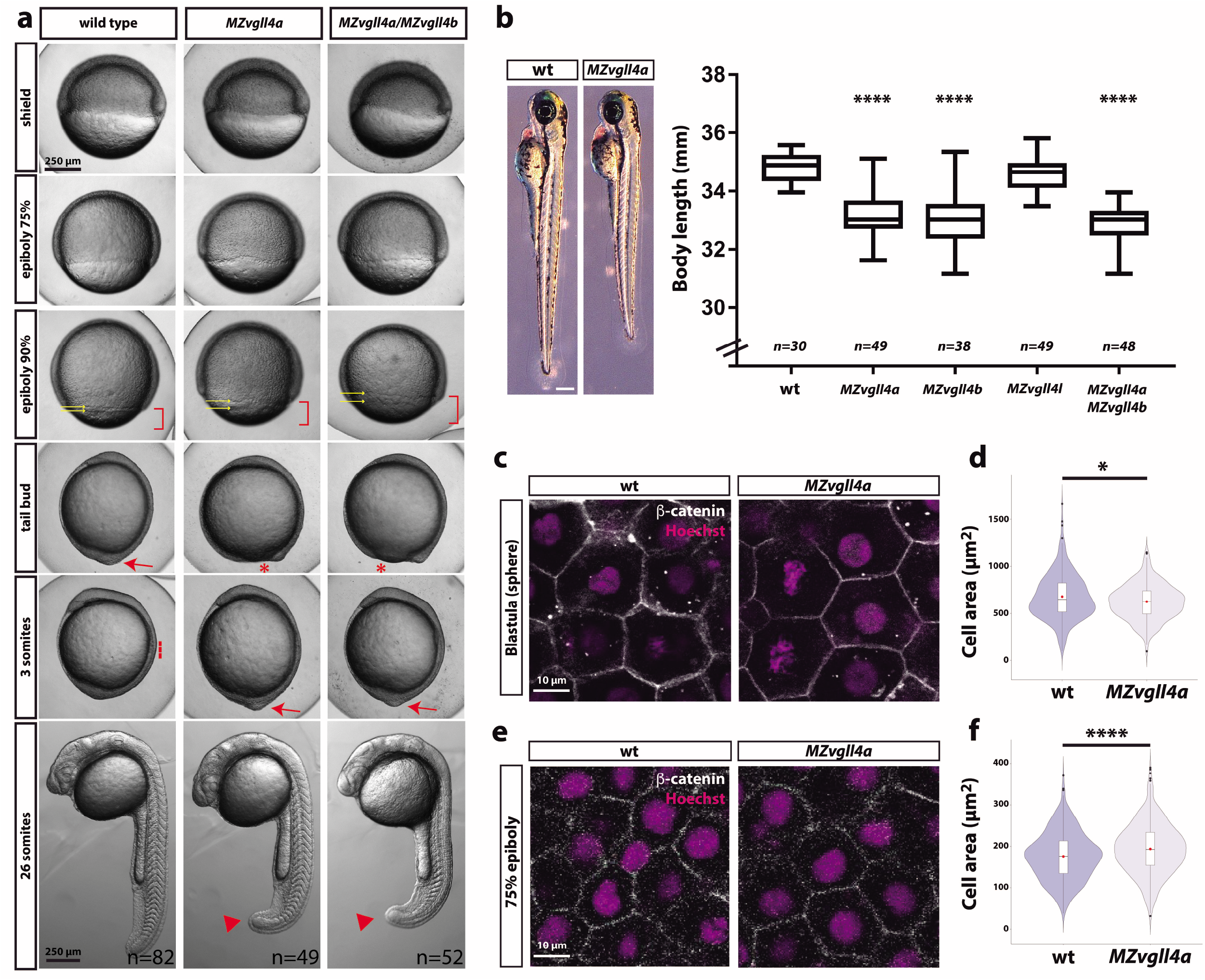
Epiboly progression in *MZvgll4* embryos is delayed. **a)** Bright field images of wt, *MZvgll4a* and *MZvgll4a/MZvgll4b* embryos at different developmental stages as indicated in the panels for wt embryos. Note the delayed epiboly (red brackets) and blastopore closure (red asterisks) as well as the increased distance between the EVL and DEL margins (yellow arrows) in *MZvgll4a* and *MZvgll4a/MZvgll4b* embryos as compared to wt. Blastopore closure (red arrow) in mutants occurs when wt embryos have initiated somitogenesis (red dotted line). At the 26 somite stage, the tail of mutants is still not fully elongated (red arrowheads). **b**) Lateral views of wt and MZ*vgll4a* larvae at 3dpf (left) and graph (right) showing the quantification of the body length from wt, *MZvgll4a, MZvgll4b, MZvgll4l* and *MZvgll4a/MZvgll4b* larvae at 3dpf. Note that MZ*vgll4a*, MZ*vgll4b* and MZ*vgll4a/vgll4b* but not MZ*vgll4l* mutants are shorter that the wt. Kruskal-Wallis test. **** p<0.0001. **c, e**) Confocal images of dorsally viewed wt and *MZvgll4a* blastulas at 75% epiboly stage, stained with β-catenin antibody and Hoechst to visualize cellular plasma membranes and nuclei, respectively. Settings for image acquisition were adjusted in each sample to visualize clearly the plasmamembranes, thus enhancing the mutant levels. **d, f**) Violin-plots of the cell area from embryos shown in c, e. Data in d represent n=252 wt and n=232 *MZvgll4a* cells from 8 and 7 blastulas, respectively. Mann-Whitney test. p= 0.0384. Data in f represent n= 431 wt and n=611 *MZvgll4a* cells from 7 and 12 embryos, respectively at 75% epiboly stage. t-test, p <0.0001.

To understand the possible reason of the *MZvgll4a* developmental delay, we stained the blastomere plasma membranes with antibodies against β-catenin. At sphere stage, the mean area of individual DEL cells in *MZvgll4a* embryos was significantly smaller than that of wt (Fig. 1c-d) but it was then significantly larger as epiboly progressed (Fig. 1e, f). Similar cell size differences have been observed in mutants with slow epiboly, such as, for example, the E-cadherin mutants (Kane *et al*, 2005).

Together these data suggest that *vgll4a* and *vgll4b* function contributes to epiboly progression and their respective absence has consequences still visible at larva stages. Notably however, maternal *vgll4a* and *vgll4b* seem to act independently and cannot compensate each other function.

### Maternal but not zygotic *vgll4a* contribution is necessary for a timely development

Many factors implicated in epiboly are maternally expressed (Lepage & Bruce, 2010). The phenotype observed in *MZvgll4a* embryos was not observed in Z*vgll4a* mutant embryos obtained from heterozygous in-crosses (data not shown), suggesting that maternal but not zygotic *vgll4a* contributes to epiboly progression.

To test this possibility, we mated wt, heterozygous and mutant *vgll4a* fishes to generate larvae of different genotypes with or without *vgll4a* maternal contribution (Fig. 2a, c). Determination of the body length of the resulting larvae showed that, similarly to *MZvll4a, Mvgll4a^+/-^* embryos had an antero-posterior axis shorter than that of *vgll4a^+/-^* and wt embryos (Fig. 2b). This indicates that zygotic *vgll4a* cannot compensate for the lack of its maternal function. To demonstrate that maternal *vgll4a* is indeed sufficient, we crossed *vgll4a* mutant males with *vgll4a* heterozygous females and *vgll4a* heterozygous males with *vgll4a* mutant females to obtain *Zvgll4a* and *MZvgll4a* embryos, with or without maternal *vgll4* contribution, respectively (Fig. 2c). Notably, the body length of the resulting *Zvgll4a* larvae at 3dpf was very comparable to that of wt (Fig. 2d), indicating that the observed developmental delay is linked exclusively to the absence of *vgll4a* maternal contribution.

**Figure 2.**
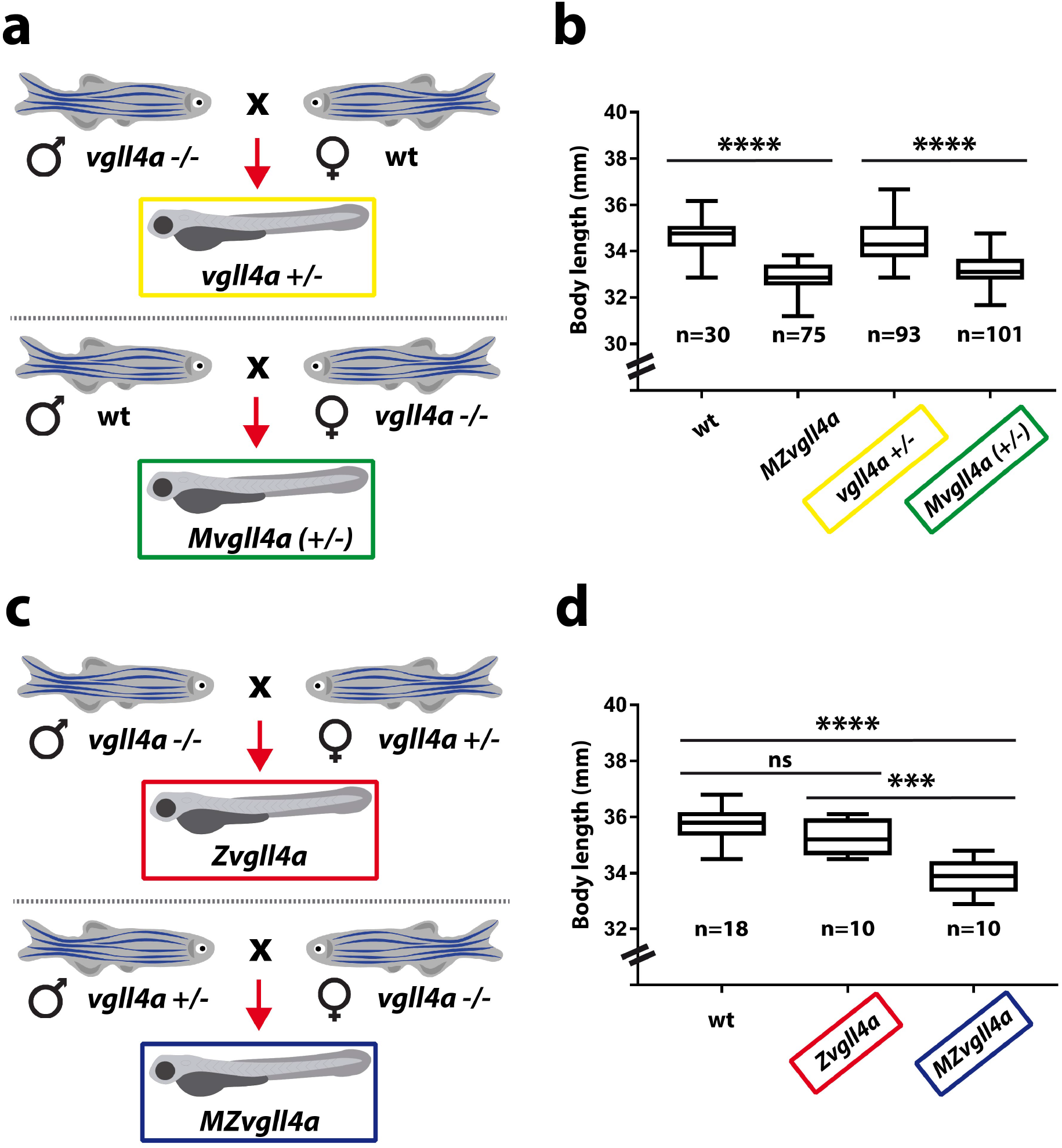
Zygotic *vgll4a* function is dispensable for timely embryonic growth. **a, c**) Schematic representation of the mating strategy used to obtain embryos of the desired genotypes with or without *vgll4a* maternal contribution. **b, d**) Box plots of the body length from 3dpf larvae of the indicated genotypes. Note the decreased body length only in embryos with absent or haplo-insufficient maternal *vgll4*a contribution as compared to wt. The number of analyzed embryos is indicated below each plot. Data in b were analyzed with Kruskal-Wallis test. **** p<0.0001, whereas those in d) with One-Way ANOVA. ns, not significant. *** p< 0.001. **** p< 0.0001.

We next asked if *vgll4b* had a similar maternally restricted function. However, whereas *MZvgll4b* are significantly shorter than wt, the body length of *Mvgll4b^+/-^* larvae was undistinguishable from that of wt (Fig. S3). This suggests that zygotic *vgll4b* can compensate for its maternal function.

To verify that the phenotype observed in *MZvgll4a* larva is directly linked to the absence of the maternally inherited mRNA/protein, we attempted to rescue *MZvgll4a* embryonic growth by injecting either the *vgll4a-HA* mRNA, its mutated version or the human VGLL4 protein in one cell stage embryos (Fig. S4a). None of the two mRNAs could rescue the growth of the mutant embryos (Fig. S4b), despite the presence of the HA-tagged protein in the blastomeres as determined by immunofluorescent staining (Fig. S4c, d). Injection of recombinant human VGLL4 protein instead partially rescued the growth of *MZvgll4a* embryos (Fig. S4e), likely because its immediate availability can compensate for the maternal component.

Taken altogether, these observations demonstrate that maternal, but not zygotic, *vgll4a* is essential for timely embryonic development.

### Vgll4a sustains yap1 signaling to promote embryonic development

Vgll4 competes with Yap1 for TEAD binding in different biological contexts, thereby antagonizing its signaling (Jiao *et al*, 2014; Koontz *et al*, 2013; Zhang *et al*, 2014). According to this mechanism, the *MZvgll4a* mutants should present an over-activation of Yap signaling and thus, Yap inhibition should rescue the *MZvgll4a* phenotype.

Verteporfin acts as an efficient inhibitor of YAP-TEAD interaction, reducing Yap1 signaling in different models (Liu-Chittenden *et al*, 2012), including the zebrafish (Fillatre *et al*, 2019; Grampa *et al*, 2016). We thus soaked wt and the *MZvgll4* embryos in verteporfin from the one cell stage up to 8hpf and thereafter let the embryos develop in fresh medium until 3 days, when their body length was measured (Fig. 3a). Wt and *MZvgll4l* body size was reduced (Fig. 3b, c) to an extent similar to that of *MZvgll4a* larvae (Fig. 1b). Notably instead, verteporfin treatment had no effect on the length of *MZvgll4a* and *MZvgll4a;MZvgll4b* larvae, which were undistinguishable from their respective untreated controls (Fig. 3d, e). This suggests that Vgll4a and Yap1 do not compete but rather pull in the same direction. To verify this possibility, we asked if Yap signaling was decreased in *MZvgll4a* embryos, using the expression levels of the Yap1 targets *ccn1* and *ccn2* as read-outs (Zhang *et al*, 2011). qPCR analysis showed that at sphere stage *ccn1* transcripts in *MZvgll4a* embryos were significantly reduced compared to wt, although this difference was no longer evident at 75% epiboly stages (Fig. 3f, g). The *ccn2a* and *ccn2b* mRNAs were undetectable at these stages (data not shown).

**Figure 3.**
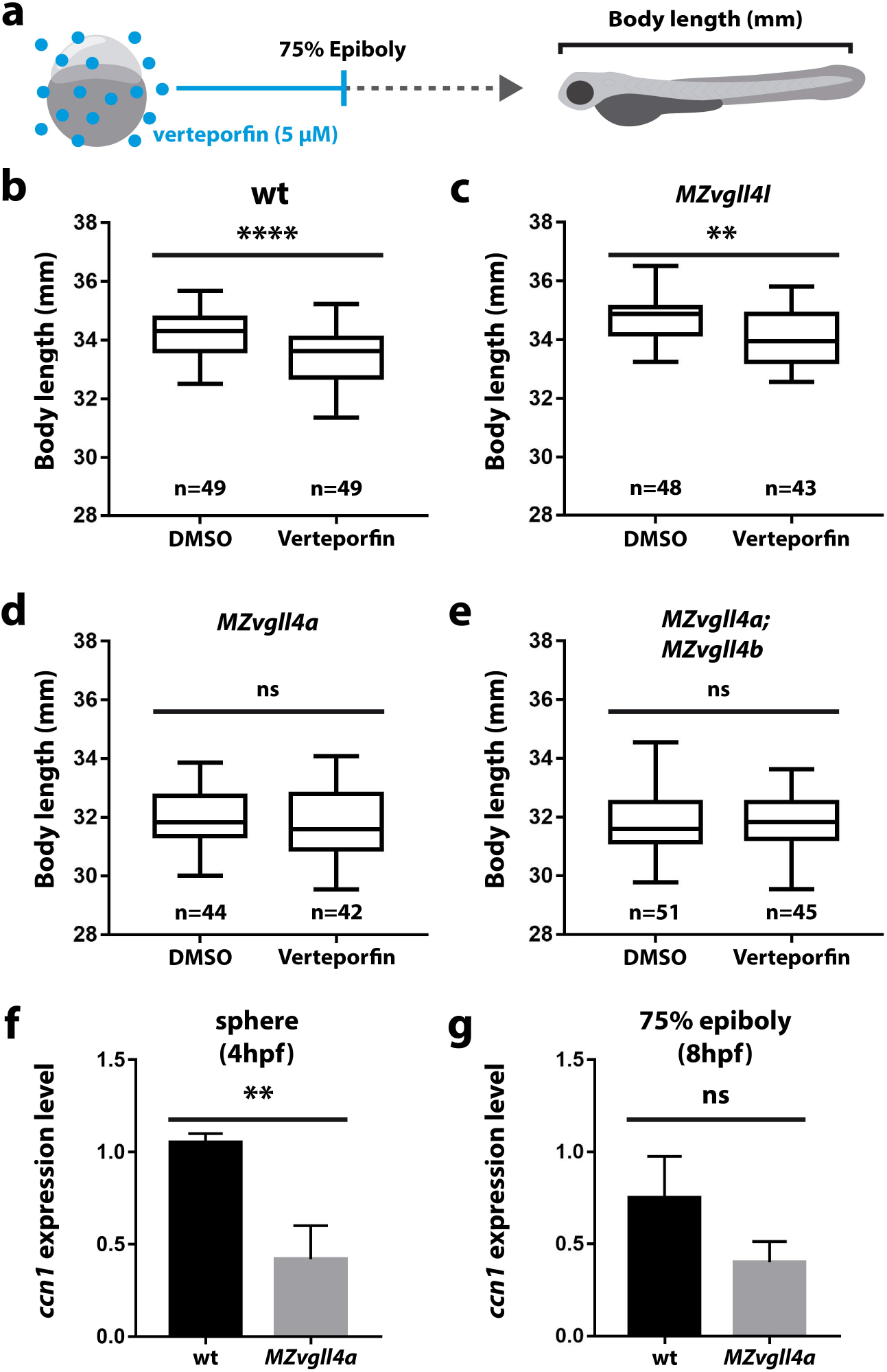
Vgll4a acts upstream of yap activity to promote embryonic growth. **a)** Schematic representation the experimental design. **b-e**) Box plots of the body length from wt (b), *MZvgll4l* (c), *MZvgll4a* (d) and *MZvgll4a;MZvgll4b* (e) embryos grown in the presence of verteporfin or DMSO. Note that the drug has no effect on the already reduced body length of *MZvgll4a* and *MZvgll4a;MZvgll4b* larvae but reduces that of wt and *MZvgll4l* larvae. Data were analyzed with Mann-Whitney test. In b, c, d and e significance is as follow: p<0.0001; p=0.0012; p=0.5706 and p=0.6064. **f, g**) The graphs show the expression level of the Yap-TEAD transcriptional target *ccn1* at sphere and 75% epiboly stage in wt and *MZvgll4a* embryos as determined by Q-RT-PCR analysis. Note the significant reduction of *ccn1* expression in the mutants. t test. p=0.0042 in f and p=0.0746 in g. ns, not significant, ** p<0.01; **** p<0.0001.

Altogether these data indicate that maternal *vgll4a* is required to sustain yap1 signaling during zebrafish gastrulation.

### Maternal *vgll4a* is required for constriction of the actomyosin ring during epiboly

Loss or knockdown of *yap* in medaka, zebrafish and *Xenopus* embryo causes a delay in blastopore closure (Gee *et al*, 2011; Porazinski *et al*, 2015). Further in the medaka fish *yap* mutant, *hirame* (*hir*), this delay is associated with a reduced actomyosin-mediated tissue tension, as also observed after the combined knockdown of *yap* and *taz* in zebrafish (Porazinski *et al*, 2015). Furthermore, in *hir* mutants the expression of the Yap signaling effector, *arhgap18*, which controls tissue tension, is downregulated (Porazinski *et al*, 2015).

We thus reasoned that if maternal vgll4a sustain yap1 signaling, the *MZvgll4a* phenotype should be also associated with alterations of the actomyosin ring and a reduction of the epibolic *arhgap18* expression. Indeed, in *MZvgll4a* embryos, the actomyosin ring was thinner (Fig. 4a-b’, e) and the DEL and EVL margins more separated than those of wt embryos (Fig. 4a-b’, f). The contour of EVL cells close to the EVL margin was also irregular with evaginations that resembled lamellipodia structures and loss of cellular contacts (Fig. 4c-d’). Furthermore, and similarly to the *hir* mutants, the transcript levels of *arhgap18* were reduced as compared to those of wt (Fig. 4g).

**Figure 4.**
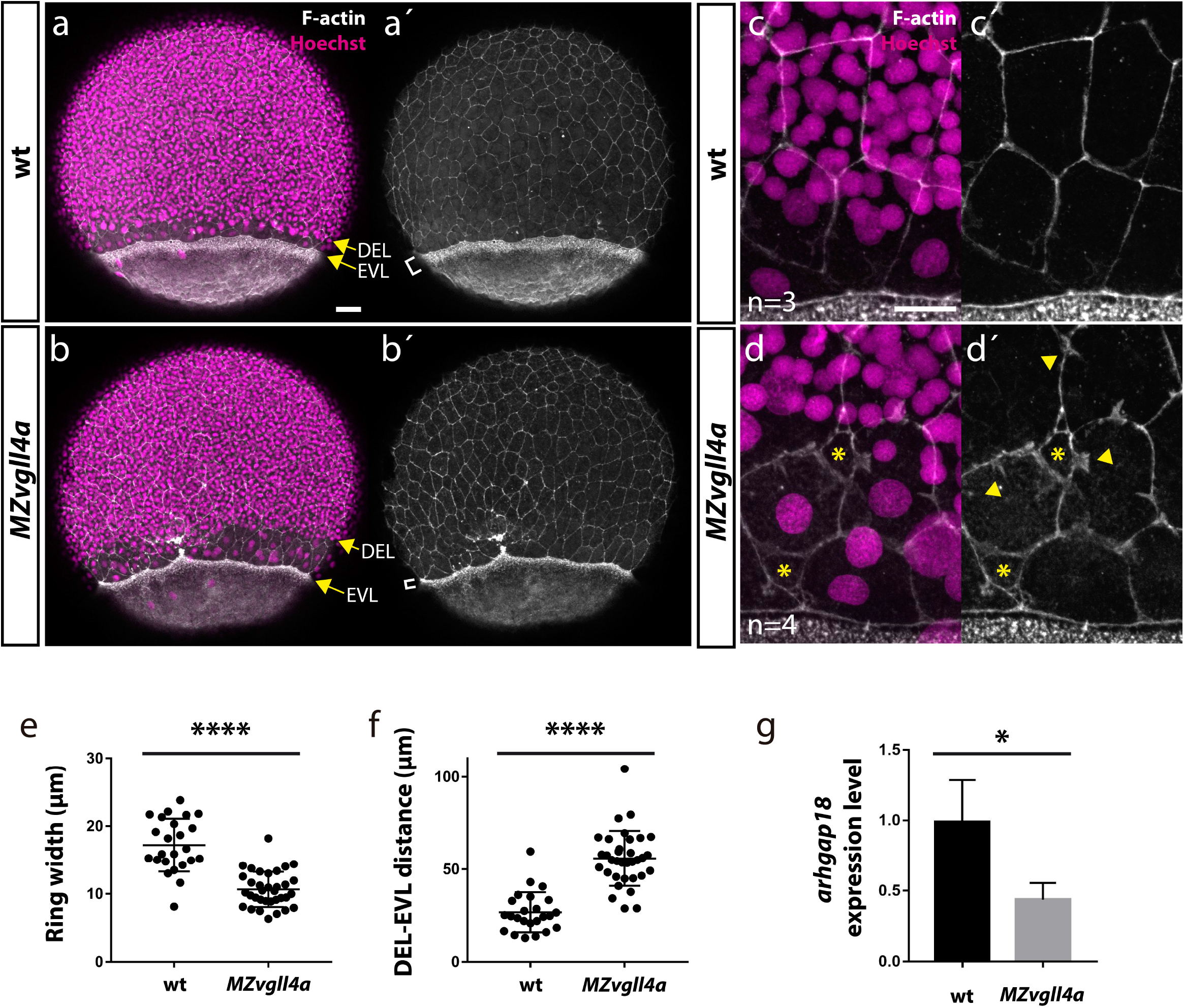
Maternal *vgll4a* is required for proper organization of the actomyosin ring. **a-d’**) Confocal images of wt and *MZvgll4a* embryos at 75% epiboly stage (lateral views) stained with phalloidin and Hoechst to visualize F-actin and nuclei, respectively. Note that the actomyosin ring in *MZvgll4a* embryos is thinner (white brackets in a’,b’) and rather separated from the DEL margin (yellow arrows in a, b) as compared to wt embryos. Note also that F-actin distribution of EVL cells in *MZvgll4a* embryos is ruffled and disorganized (yellow arrowheads in d’) in contrast to the well aligned distribution in wt embryos. Asterisk in d, d’ indicates loss of cell-cell contacts. **e, f**) Quantification of the actomyosin ring width and DEL-EVL distance in wt (n=24) and *MZvgll4a* (n=34) embryos (Mann-Whitney test. ****, p<0.0001). **g**) The graphs show the expression level of *arhgap18* transcripts in *MZvgll4a* and wt embryos at 75% epiboly stage (t-test. p=0.041), as determined by qRT-PCR analysis. *, p<0.05. Scale Bars, a-b’ 50μm, c-d’, 20μm.

Taken together these observations support the idea that vgll4a function is a requisite for yap1-dependent actomyosin ring contractibility and thus epiboly progression.

### Maternal *vgll4a* promotes E-cadherin/β-catenin distribution at the blastomere plasma membrane

Cells probe tension through plasma membrane proteins and then transmit the information to mechano-sensors such as Yap1 by means of cytoskeletal rearrangements (Panciera *et al*, 2017). Given that maternal vgll4a seemed to act upstream of yap1, we hypothesized that its activity could regulate blastomere adhesion and thus their tension probing capacity. Indeed, in cancer cells VGLL4 regulates the transcription of E-cadherin (Li *et al*, 2015; Song *et al*, 2019) and the E-cadherin/α-Catenin/β-Catenin adhesion complex is an upstream regulator of Yap1 in different biological contexts (Kim *et al*, 2011; Schlegelmilch *et al*, 2011; Silvis *et al*, 2011). Furthermore, the epibolic phenotype of *MZvgll4a* embryos resembled that of the pou5fl/Oct4 deficient *MZspg* embryos (Lachnit *et al*, 2008; Song *et al*, 2013). In these mutants, deep cells move with a considerable delay in relation to the actin-depleted EVL margin, EVL cells form an abnormal number of lamellipodia (Lachnit *et al*, 2008; Song *et al*, 2013) and present a defective E-cadherin endosomal trafficking in blastomeres (Song *et al*, 2013). We thus compared the distribution of F-actin, β-catenin and E-Cadherin in wt and *MZvgll4a* embryos, as read outs of blastomere cohesion (Yap *et al*, 2018).

F-actin, β-catenin and, to a lesser extent, E-cadherin signal intensity of *MZvgll4a* EVL cells was significantly decreased at sphere stage (Fig. 5a-h). These changes were less evident in DEL cells, where only β-catenin was significantly reduced at sphere (Fig. 5i-p) and shield (Fig. S5) stages. Notably, the overall E-cadherin mRNA expression levels (*cdh1*) in wt and *MZvgll4a* embryos were similar at both sphere and shield stage (Fig. S6a, c). β-catenin acts as an effector of Wnt signaling and thus its decreased levels at the plasma membrane could be a consequence of an over-activation of the pathway. To test this possibility, we determined the expression levels of different Wnt targets and read-outs. No difference was observed in the levels of *axin1, axin2* and *lef1* expression between *MZvgll4a* and wt embryos at both sphere and shied stage (Fig. S6b, d).

**Figure 5.**
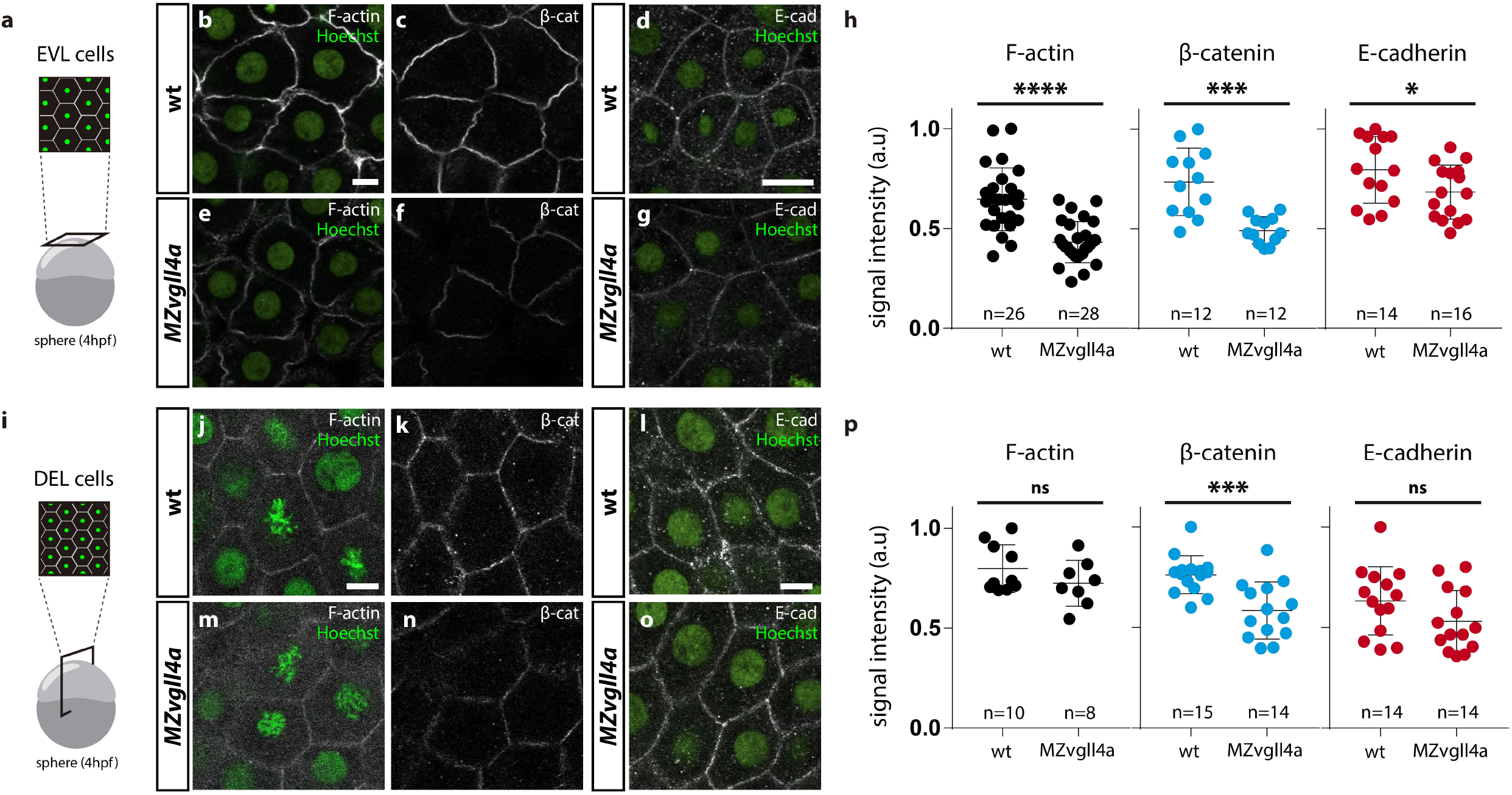
Maternal vgll4a is required for plasma membrane localization of the E-cadherin/β-catenin complex. **a, i)** Schematic representation of the different imaging strategies. **b-g; j-o)** Confocal images of F-actin (b, e, j, m), β-catenin (c, g, k, n) and E-cadherin (d, g, l, o) distribution in EVL (b-g) and DEL (j-o) cells in wt and *MZvgll4a* embryos at sphere stage. Embryos were counterstained with Hoechst (nuclei, green). **h, p**) The graphs depict the fluorescent signal intensity (in arbitrary units, a.u) for F-actin, β-catenin and E-cadherin in EVL (h) or DEL (p) cells of wt and *MZvgll4a* embryos (Mann Whitney test. h) E-cadherin; p= 0.0425. p) F-actin, p=0.3154; β-catenin, p=0.0005 and E-cadherin, p=0.1087. ns, not significant, * p<0.05; *** p<0.001; **** p<0.0001. Scale bar, 10μm.

All in all, these data indicate that maternal vgll4a promotes E-cadherin/β-catenin localization at the blastomeres’ plasma membrane and hence their actin cortex distribution. Blastomere cohesion, in turn, enables yap1-mediated mechano-transduction and actomyosin ring constriction driving epiboly progression.

## Discussion

The genome of a fertilized egg is transcriptionally inactive. Thus, the RNAs and proteins deposited in the eggs by the mother are responsible for coordinating the first morphogenetic events that takes place during gastrulation, initially single-handed and then in cooperation with the increasingly available zygotic gene products (Solnica-Krezel, 2020). In the zebrafish egg, these maternally derived molecules represent a large proportion of all possible gene products (Harvey *et al*, 2013). However, how many of them are critical for epiboly initiation and progression is still poorly defined. Genome wide (Kane *et al*, 1996) and specific maternal-effect mutational screens in zebrafish have identified only a limited number of mutants with an epiboly phenotype (Wagner *et al*, 2004; Dosch *et al*, 2004) and their subsequent characterization supported an important contribution of adhesion molecules and cytoskeletal components (Bruce, 2016). Other studies have thereafter identified a few maternally inherited transcriptional regulators, including eomes, foxh1, pou5fl/oct4, nanog and yap1 (Bruce *et al*, 2005; Reim & Brand, 2006; Pei *et al*, 2007; Lachnit *et al*, 2008; Song *et al*, 2013; Porazinski *et al*, 2015; Veil *et al*, 2018; Gagnon *et al*, 2018). In this still fragmented scenario, our study adds a new component to the genetic network coordinating zebrafish epiboly, showing an important role for maternally inherited vgll4a, and, in part, for its paralog vgll4b. Notably, it also shows that vgll4a is required to support, rather than antagonise, yap1 signalling and to coordinate blastomere adhesion/cohesion. Our data further confirm that this cohesion is a fundamental piece of the self-sustained bio-mechanical regulatory loop underlying epibolic morphogenetic rearrangements, which culminates with blastopore closure at the end of gastrulation.

Zebrafish epiboly can be divided in two phases: initiation or doming and progression. As a limitation, our study does not clearly define the spatio-temporal window in which vgll4a activity is required. However, the large majority of factors implicated in doming are maternally expressed, as both *vgll4a* and *vgll4b* are (Xue *et al*, 2018). This may place their activity at epiboly initiation even if the phenotypic consequences of their inactivation are observed only later. Such a time-lag has been indeed reported for other factors contributing to epiboly initiation (Lachnit *et al*, 2008; Sun *et al*, 2017). A recent study has shown that at sphere stage the central blastula becomes “fluid” as a consequence of loss of cell-cell adhesion and increased cell division (Petridou *et al*, 2019). The blastoderm margins instead maintain cell-cell adhesion and thus tension thanks to the activity of Wnt11 non-canonical signalling (Petridou *et al*, 2019), known to control E-cadherin availability at the plasma membrane (Ulrich *et al*, 2005). If this differential fluid vs tense state is perturbed, the ability of the blastula to react to mechanical forces is altered and epiboly becomes defective (Petridou *et al*, 2019). The phenotype of *MZvgll4a* mutants could be explained within this frame. We show that *vgll4a* role in epiboly can be considered as strictly maternal (Solnica-Krezel, 2020) and thus could likely act upstream of Wnt11 signalling. Vgll4a-mediated control of Wnt signalling will impinge on E-cadherin/β-catenin availability at the plasma membrane and thus differential blastoderm viscosity critical for epiboly progression. In support of this possibility, *vgll4* paralogs have been shown to control the expression of non-canonical Wnt signaling components, including *wnt11*, *fzd8a* and *fzd10* (Fillatre *et al*, 2019). Alternatively, *vgll4a* could interfere with non-canonical Wnt signaling indirectly through the regulation of Wnt/β-catenin canonical pathway, as the two branches of the pathway have been previously shown to have antagonistic effects in morphogenesis (Cavodeassi *et al*, 2005). Indeed, VGLL4 seems to interfere with the formation of a TEAD4-TCF4 complex, which promotes cell proliferation in colorectal cancer (Jiao *et al*, 2017). Furthermore, VGLL4 overexpression suppresses nuclear β-catenin levels and inhibits migration and invasion of gastric cancer cells, while its inactivation has opposite effects (Li *et al*, 2015). As an additional possibility, vgll4a could directly regulate E-cadherin expression as observed in both gastric and breast cancers (Li *et al*, 2015; Song *et al*, 2019), thereby influencing the membrane levels of E-cadherin/β-catenin complexes. According to our data the latter possibilities are however less likely. We observed a small, although significant, decrease of E-cadherin membrane localization only in EVL cells of *MZvgll4a* embryos at sphere stage, and no significative changes in the mRNA levels of *cdh1* or in those of three read-out of Wnt/β-catenin signalling. However, our analysis was performed using the entire embryos, precluding the identification of changes occurring only in a small subset of the cells. Furthermore, we cannot rule out that the *MZvgll4a* phenotype might be associated with E-cadherin destabilization or alteration of its trafficking, as reported for the zebrafish *wnt11/slb* and *oct4/MZspg* mutants (Song *et al*, 2013; Petridou *et al*, 2019).

Although our study focuses on *vgll4a*, we report that the related *vgll4l* has no role in epiboly, consistent with its lack of early expression (Xue *et al*, 2018) and its suggested functional diversification (Fillatre *et al*, 2019). On the contrary, we show both *vgll4a* and *vgll4b* seem to be important for oocyte fecundity and that *MZvgll4b* and *MZvgll4a* mutants present a very comparable developmental delay. This indicates that *vgll4b* has also an important role in both oocyte and gastrulation. However, in contrast to what observed for *vgll4a*, maternal *vgll4b* activity can be compensated by its zygotic counterpart. Furthermore, *MZvgll4a;MZvgll4b* double mutants are phenotypically very comparable to the two single mutants, suggesting that the two paralogs do not compensate each other function and perhaps act in a different spatio-temporal window that converges in the same shorter axis phenotype. At later stages of development, *vgll4b* has been implicated in erythropoiesis, terminal differentiation (Wang *et al*, 2020), heart valvulogenesis (Xue *et al*, 2019) and establishment of the left-right asymmetry in coordination with *vgll4l* (Fillatre *et al*, 2019). Our *vgll4a* mutants do not show traits linked to similar functions, at least in a gross morphological analysis, further supporting a possible diversification of the paralog activity.

Vgll4 has been initially described as co-transcriptional repressor that interacts with Tead transcription factors, thereby acting as an as antagonist of Yap signalling (Guo *et al*, 2013; Zhang *et al*, 2014). This function has been thereafter validated in multiple contexts especially in tumour development (Yamaguchi, 2020). Our study not only shows that maternal vgll4a is required for epiboly but also that it does so supporting rather than antagonising yap1 signalling. The generation of a transgenic Yap/Taz-Tead reporter zebrafish line faithfully highlights pathway activation from 22 hpf (Miesfeld & Link, 2014). Using this same line we have been unable to detect reporter expression at epiboly stages and have thus used alternative approaches to determine yap1 signalling, including the expression of the yap1 downstream target *ccn1* and embryonic treatment with verteroporfin. These approaches show a decreased yap1 signalling in *MZvgll4a* mutants and support a possible role of vgll4a upstream of yap1, as we have proposed above. However, our analysis has been performed at the scale of the whole embryo, taking epiboly progression, actomyosin organization and cell cohesion as read-out of vgll4a function in morphogenetic rearrangement. These rearrangements however highly depend on mechanical and biochemical events at the cellular and subcellular scale (Stooke-Vaughan & Campàs, 2018; Petridou *et al*, 2017). In this scenario, we cannot exclude that the delay in epiboly progression observed in *MZvgll4a* embryos represents the macroscopic result of a series of mechano-chemical feedback loops at the cellular scale (Hannezo & Heisenberg, 2019), involving both vgll4a and yap1 in the same or even different cell populations.

Nevertheless, and independently of the relative position of vgll4a and yap1 in epiboly regulation, our data clearly show that vgll4a is required for a proper balance between tissue tension/cohesion and contractibility, two factors that contribute to mechanical stress (Wozniak & Chen, 2009). Thus, in the absence of vgll4a, cellular cohesion mediated by adhesion complexes is poor, impairing embryonic mechanosensation (Fig. 6). This impairment decreases yap-dependent mechano-transduction, which, in turn, affects the contraction of the actomyosin network, delaying epiboly progression (Fig. 6).

**Figure 6.**
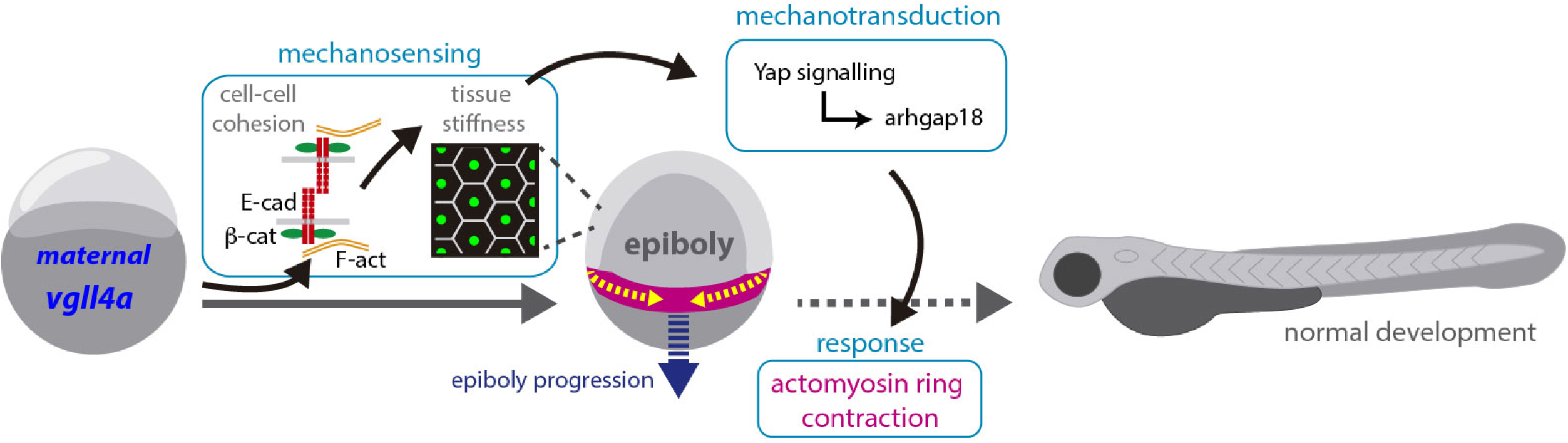
Proposed model for maternal vgll4a contribution to zebrafish epiboly progression. Maternal *vgll4a* promotes plasma membrane localization of the E-cadherin/β-catenin complex in the amount required for an adequate cohesion among blastomeres. This cohesion threshold allows tissue mechano-sensing and thus yap1-dependent mechano-transduction. Signal transduction impacts on the organization and function of the actomyosin ring, thereby promoting timely epiboly progression. E-cad, E-cadherin; β-cat, β-catenin; F-act, F-actin.

In a broader context, our data suggest that up-regulation of Vgll4 expression may serve to enhance the mechano-sensing properties of some tissues, perhaps restoring an unbalanced back-and-forth dialogue between biochemical and mechanical cues, which has been described in pathological conditions, including different type of primary and metastatic cancers (Deng & Fang, 2018).

## Materials and methods

### Maintenance of fish lines

Adult AB/TUE wild type and mutant (*vgll4a, vgll4b, vgll4l*) zebrafish were maintained at 28°C on 14/10 h light/dark cycle. Embryos were raised at 28°C and staged according to the hours post fertilization (hpf) and their morphology. Embryos were growth in E3 medium (NaCl, 5 mM; KCl, 0.17 mM; CaCl2, 0.33 mM; MgSO4, 0.33 mM; 5,10% Methylene Blue). The ethical committee for Animal Experimentation of the Consejo Superior de Investigaciones Científicas (CSIC) and of the Comunidad Autónoma de Madrid approved the procedures used in the study.

### Zebrafish mutants generation

Zebrafish mutant lines were generated using CRISPR/Cas9 technology. The gRNA were designed using the CHOP-CHOP tool (Labun *et al*, 2019) looking for potential disruption of enzyme restriction sites. gRNAs were synthesized using PCR templates as described in (Varshney *et al*, 2016). Cas9 protein (300 ng/μL; EnGen^®^ Spy Cas9 NLS, New England Biolabs) and gRNAs (100 ng/μl; Table S1) were co-injected into one-cell stage zebrafish embryos. F0 embryos were raised and outcrossed with AB/TUE wt. PCR amplification on genomic DNA isolated from tail clips of F1 zebrafish embryos was performed to identify disruption of specific restriction sites (Table S2). The DNA of potential mutants was thereafter sequenced and the selected embryos were raised to adulthood. We selected mutants in which the reading frame was disrupted and truncated as follow: *vgll4a*-S85Mfs15, *vgll4b*-P156Rfs5 and *vgll4l* P111Hfs61 (see Fig. S1 for more details).

### Embryo injections

Zebrafish embryos were injected at the one cell stage using a IM-300/Narishige microinjector. Recombinant humanVGLL4 protein (0.19 ug/μl; Abnova, Ref: H00009686-P01) and *vgll4a*-HA mRNA (100 ng/μl) were used in rescue experiments. Phosphate buffer saline (PBS) containing 0.1% BSA or mutated *vgll4a*-HA mRNA (cloned from the *vgll4a*-S85Mfs15 line) were injected as controls.

### Tissue processing and immunochemistry

Embryos were fixed by immersion in 4% paraformaldehyde in 0.1M phosphate buffer pH 7.2 (wt/vol) overnight at 4 °C. Embryos were then washed in PBS with 0.5% Triton-X-100, incubated in a 15% sucrose-PBS solution (wt/vol), embedded and frozen in a 7.5% gelatin in 15% sucrose solution (wt/vol). Cryostat sections or whole embryos were stained using standard protocols and antibodies against the following antigens: HA (1:250, Sigma), β-catenin (1:300, BD Bioscience) and E-cadherin (1:300, BD Bioscience). Incubation with phalloidin (1:200, Sigma) was used to detect actin. Incubation with appropriate secondary antibodies was performed following standard procedures.

### *vgll4a-HA* cloning

*vgll4a* was amplified from cDNA of wt or *MZvgll4a* (as control) embryos using the following primers (Fw: 5’-GGAATCAACAGTTAGCGTGCT-3’; Rv: 5’-aaCTCGAGTCAAGCGTAATCTGGAACATCGTATGGGTAAGACTGACC AACATGATTG-3’). The amplicon, which includes the hemagglutinin (HA) epitope, was cloned by TA-cloning in the pSC-A vector using the StrataClone PCR Cloning Kit (Agilent). The *vgll4a-HA* fragment was thereafter obtained from the pCS-A construct using EcoRI/XhoI digestion and cloned in the pCS2 vector. The pCS2 construct, linearized with NotI, was used to synthesize mRNA from the wt or mutated *vgll4a* version, tagged with HA, using the *mMESSAGE mMACHINE SP6 Transcription kit* (Invitrogene) following manufacturer’s instructions. After transcription, mRNAs were purified using the NucleoSpin^®^ RNA Clean-up kit (Machery Nagel).

### Quantitative RT-PCR analysis

Total RNA was isolated from wt and mutant embryos (n=30, for each genotype) using TRIzol (Sigma) according to the manufacturer’s instruction. Each experiment was performed with biological triplicates. 5 μg of total RNA was used to synthesize the first-strand cDNA using the First-Strand cDNA synthesis kit (GE Healthcare) with a pd(N)_6_ primer. Each quantitative RT-qPCR reaction was performed using the GoTaq qPCR Master Mix kit (Promega). For a 10 μl reaction, 4 μl of cDNA (2.5 ng/μl) was mixed with 1 μl of primers (2.5 μM; Table S3) and 4 μl master mix. Reaction was incubated at 95°C for 10 min, then at 95°C for 15 s and 40 cycles and at 60°C for 60 s. The levels of the *eef1a1l1* mRNA were used as housekeeping reference (Xu *et al*, 2016).

### Pharmacological treatment

Embryos were incubated at 28°C from the one cell stage to 75% epiboly (8 hpf) in E3 medium containing 5 μM of the Yap1 antagonist verteporfin (5305, Tocris) dissolved in DMSO or the same volume of DMSO as control.

### Imaging

Sections were analysed with DM or confocal microscope. Zebrafish embryos at different stages were observed and photographed using a stereomicroscope and DFC500, DFC350 FX cameras (Leica Microsystems). For sections or whole embryo staining, LSM710 confocal laser scanning microscope coupled to an AxioObserver inverted microscope (Zeiss) were used to obtain digital images, which were then processed and analysed with ImageJ (Fiji) software. Images shown in Figures 1, 4, 5, S1, S2, S4 and S6 were assembled using the Photoshop CS5 software.

### Quantification and statistical analysis

All quantifications were performed using the ImageJ (Fiji) software. 72hpf embryonic larvae were treated with tricaine and photographed in lateral views at 13 X magnification to determine their body length. The area of individual and centrally positioned blastoderm cells from wt and mutant embryos was determined by tracing their perimeter on confocal images of whole embryos stained with β-catenin antibody. Only one image per embryo was used. The position and width of the actomyosin ring was determined on z-stack files obtained from the confocal images of whole embryos. The z-stack files were used to generate maximum intensity projections. The mid-region of the actomyosin ring and DEL/EVL margins were positioned in the centre of the image to avoid possible visual distortions due to the embryo curvature and the region of interest (ROI) was selected. Ten different measurements of the width of the actomyosin ring as well as the distance between DEL-EVL were obtained along the ROI using the “straight” selection tool. The ten measurement *per* embryo were averaged to obtain a more accurate value and their mean was used in the comparative analysis. Immunofluorescence intensity quantifications were performed using mid-plane of z-stack files obtained from the confocal imaging of β-catenin, F-actin and E-cadherin staining on whole embryos (EVL cells) or cryostat sections of different embryos (DEL cells). A rectangular ROI was used to measure the mean gray value of the selected image as indicator of signal intensity. GraphPad Prism 7 statistic software was used to analyze the data using t-test for two groups with parametric distribution and with Mann-Whitney test for two groups with no parametric distribution. One-way ANOVA test was used for more than two groups with parametric distribution and Kruskal-Wallis test for no parametric distribution. Statistical difference between pools of treated and control embryos was determined with the Pearson’s X2 test. Violin-plots in Figure 1 were obtained with the R-software and the ggplot2 package. Error bars indicate s.e.m. in all graphs.

## Acknowledgements

We wish to acknowledge the excellent technical assistance of Alfonso Gutierrez Garcia in the fish facilities and the staff of the CBMSO Image analysis and Genomic and Next Generation Sequencing facilities. This work was supported by grants from the Spanish MINECO (BFU2016-75412-R with FEDER funds); AEI (PID2019-104186RB-I00 and RED2018-102553-T) and a grant from the Fundación Ramon Areces to PB. CCM and MJC are supported by predoctoral contract from the CIBERER and a Juan de la Cierva postdoctoral contract from the AEI (IJCI-2016-27683), respectively. A CBM Institutional grant from the Fundación Ramon Areces is also acknowledged.

## Author contributions

CCM, MJC and PB conceptualized and designed the research study. CCM performed most of the experiments, acquired and analyzed the data. NT generated data reported in Fig. 1, 5 and Fig. S4, S6 and MJC those in Fig. 1, 4, 5 and Fig. S5. PB obtained financial support. CCM, MJC and PB wrote the paper. All authors read and approved the manuscript.

